# Species delimitation and biogeography of the gnatcatchers and gnatwrens (Aves: Polioptilidae)

**DOI:** 10.1101/271494

**Authors:** Brian Tilston Smith, Robert W. Bryson, William M. Mauck, Jaime Chaves, Mark B. Robbins, Alexandre Aleixo, John Klicka

**Author notes:** Current Address: New York Genome Center, New York, NY 10013, USA. **Corresponding Author:** Brian Tilston Smith, Department of Ornithology, American Museum of Natural History, Central Park West at 79th Street, New York, New York 10024, USA, **Email address:**.

## Abstract

The New World avian family Polioptilidae (gnatcatchers and gnatwrens) is distributed from Argentina to Canada and includes 15 species and more than 60 subspecies. No study to date has evaluated phylogenetic relationships within this family and the historical pattern of diversification within the group remains unknown. Moreover, species limits, particularly in widespread taxa that show geographic variation, remain unclear. In this study, we delimited species and estimated phylogenetic relationships using multilocus data for the entire family. We then used the inferred diversity along with alternative taxonomic classification schemes to evaluate how lumping and splitting of both taxa and geographical areas influenced biogeographic inference. Species-tree analyses grouped Polioptilidae into four main clades: *Microbates*, *Ramphocaenus*, a *Polioptila guianensis* complex, and the remaining members of *Polioptila*. *Ramphocaenus melanurus* was sister to the clade containing *M. cinereiventris* and *M. collaris*, which formed a clade sister to all species within *Polioptila*. *Polioptila* was composed of two clades, the first of which included the *P. guianensis* complex; the other contained all remaining species in the genus. Using multispecies coalescent modeling, we inferred a more than 3-fold increase in species diversity, of which 87% represent currently recognized species or subspecies. Much of this diversity corresponded to subspecies that occur in the Neotropics. We identified three polyphyletic species, and delimited 4–6 previously undescribed candidate taxa. Probabilistic modeling of geographic ranges on the species tree indicated that the family likely had an ancestral origin in South America, with all three genera independently colonizing North America. Support for this hypothesis, however, was sensitive to the taxonomic classification scheme used and the number of geographical areas allowed. Our study proposes the first phylogenetic hypothesis for Polioptilidae and provides genealogical support for the reclassification of species limits. Species limits and the resolution of geographical areas that taxa inhabit influence the inferred spatial diversification history.

## INTRODUCTION

High-throughput DNA sequencing and thorough geographic sampling within currently recognized species has led to redefining of species limits in many groups (Fujita *et al.*, 2012). This phenomenon is true for both traditionally understudied groups, such as invertebrates (e.g., Satler *et al.*, 2013), and more frequently studied groups, such as vertebrates (e.g., Leaché & Fujita, 2010). Taxonomic instability is often associated with taxa that occur in high-diversity tropical latitudes, due in part to proportionally fewer systematists working in these regions (Tobias *et al.*, 2008) and the logistical challenges working in remote areas. Although this problem is recognized (Tobias *et al.*, 2008, Gill, 2014), how best to update taxonomy given the immediate need of accurately delimited species for both applied and basic purposes, continues to be debated (Peterson & Navarro-Sigüenza, 1999, Tobias *et al.*, 2010, Remsen, 2016). Because species are the fundamental unit in biology, the correct identification of species is essential for research topics ranging from identifying conservation priority areas to inferring evolutionary history (Fujita *et al.*, 2012).

Debate over species concepts has been central to the modern and ongoing reclassification of avian taxa. Among ornithologists, the debate has centered on preferentially weighting reproductive isolation (Mayr, 1942) versus diagnosability (Eldredge & Cracraft, 1980; Cracraft, 1983) as the primary defining criterion. The strengths and weaknesses of these approaches have been thoroughly discussed in previous publications (Zink & McKitrick, 1995; Johnson *et al.*, 1999; de Queiroz, 2007). To summarize briefly, assessing whether or not closely related allopatric bird populations have achieved reproductive isolation is predominantly done by evaluating phenotypic differences and more recently, differentiation in song, often in concert with genetic data (Johnson *et al.*, 1999; Remsen, 2005; Gill, 2014). Behavioral response to song is typically not determined experimentally, but rather, taxonomic decisions are based on whether song differences are quantifiably above some threshold value (Gill, 2014). This approach requires quantifying how songs vary among and within species, and assumes that birds will perceive these similarities or differences in a predictable way. An incomplete understanding of subspecific song diversity has led to the lumping of allopatric taxa as single species (Remsen, 2005; Zink, 2006; Peterson & Navarro-Sigüenza, 2006; Gill, 2014). In contrast, phylogenetic species are delimited by identifying fixed differences in genetic or morphological characters. This approach works well for identifying independently evolving taxa under scenarios where alleles have sorted and morphological evolution is concordant with population history. However, when many loci are sampled it becomes unlikely that all gene trees will be reciprocally monophyletic among populations (Hudson & Coyne, 2002; Hudson & Turelli, 2003), making it challenging to delimit taxa with shallow evolutionary histories or gene flow after divergence. Fixed phenotypic differences also have limitations (for both approaches) because plumage evolution often does not track phylogeny (e.g., Omland & Lanyon, 2000; DaCosta & Klicka, 2008). Overall, irrespective of which of these concepts is invoked, diversity may be underestimated and/or subjectively characterized.

An alternative means of identifying species from genetic data is through the application of coalescent-based models (Fujita *et al.* 2012). These methods are not directly linked to a particular species concept, but may assume reproductive isolation in the absence of gene flow, and are used to determine if lineages are evolving independently. Among the benefits of a coalescent-based approach to species delimitation is that species limits can be tested in a probabilistic framework (Fujita *et al.*, 2012). Importantly, these models relax the assumption of reciprocally monophyletic groups and allow for the use of multilocus data, where individual gene trees are often unsorted among closely related taxa. Though these approaches are still based on simplistic models, existing models are robust to low levels of gene flow (Yang and Rannala 2010). More recently published methods do not require phylogenetic relationships among species to be known (Yang and Rannala 2014) and can incorporate phenotypic data (Solís-Lemus *et al.*, 2015). Applying model-based species delimitation to numerous taxonomic groups has shown both concordance and discordance with currently accepted taxonomies (Pons *et al.*, 2006; Leaché & Fujita, 2010; Lumbsch & Leavitt, 2011). The ability of coalescent modeling to objectively identify independently evolving lineages also provides a means of increasing the resolution in downstream analyses of evolutionary history.

Poorly resolved species limits, for example, may confound the inference of ancestral areas and spatial diversification histories. In historical biogeographic analyses that reconstruct or estimate the probability of an ancestral area (Ree & Smith, 2008; Matzke, 2014; Yu *et al.*, 2015), taxa are assigned to geographical areas and these assignments are influenced by how “lumped” or “split” a taxon is. Lumped taxa are more likely to be polytypic, geographically widespread, and found in multiple biogeographic areas than finely split taxa. Another potentially confounding factor is that the geographic resolution of an area can also be lumped or split hierarchically. A taxon can be assigned to a continental-scale area such as South America or to a more fine-scale endemic area like the Napo region of the Amazon Basin. The impact of lumping and splitting of both taxa and areas on biogeographic inference has been poorly explored, but one expectation is that lumping will provide relatively coarse-scale resolution of when and where divergence events occurred. The direction of bias would depend on how much area assignments change between lumped and split taxa, and whether divergence events of interest occurred within species. For widely distributed clades and species that span diverse geographic areas and continents, unresolved species limits could confound biogeographic inference.

In this study, we examined the evolutionary history of a geographically widespread family of New World birds, Polioptilidae. This family consists of the gnatcatchers and gnatwrens, which form three genera of diminutive insectivorous birds distributed from southeastern Canada to Argentina. *Polioptila*, the most diverse and widely distributed genus, consists of 12 species that occur in a variety of habitats including temperate forests, deserts and lowland humid forests, including one species (*Polioptila caerulea*) that undergoes long-distance seasonal migration. The other two genera, *Ramphocaenus* and *Microbates*, occur only in tropical lowland forests. *Ramphocaenus* and *Microbates* were previously placed in the distantly related suboscine family Formicariidae (Ridgway, 1911; Cory & Hellmayr, 1924), and were subsequently moved into the Old World warblers (Sylviidae) based on the presence of an oscine aftershaft (Miller, 1924) and syrinx (Wetmore, 1943). *Polioptila*, *Ramphocaenus*, and *Microbates* were collectively moved to the tribe Polioptilini forming a clade composed of all “New World slyviids” (Mayr & Amadon, 1951), but a comparison of nest characteristics challenged the notion of *Ramphocaenus, Microbates*, and *Polioptila* having a shared ancestry (Kiff, 1977). DNA hybridization and molecular sequence data later placed the Polioptilini outside of Sylviidae and sister to the New World wrens (Troglodytidae; Sibley & Ahlquist, 1990; Barker *et al.*, 2004). Further phylogenetic studies using molecular data within the family have focused on defining species limits and phylogeographic diversity in select species or complexes (Zink & Blackwell, 1998; Zink *et al.*, 2000; Zink *et al.*, 2013; Naka *et al.*, 2012; Smith *et al.*, 2012), and no study to date has examined phylogenetic relationships across all gnatcatchers and gnatwrens.

We collected multilocus data from species in Polioptilidae to estimate a species tree, delimit species using coalescent modeling, and evaluate how different taxonomic classification schemes influence biogeographic inference for the family. Our sampling focused on maximizing the representation of morphological and genetic diversity that exists within each species to identify all independently evolving lineages, regardless of current taxonomic rank. We predicted that by testing species limits using all identified lineages and coalescent modeling, we would capture an estimate of species diversity that reflects evolutionary history and demonstrate that currently recognized species either underestimate or mischaracterize species-level diversity within the family. We evaluated the implications of alternative taxonomic limits by modeling ancestral range evolution on four alternative species trees and we examined the importance of correctly defining “area” by evaluating each tree in the context of three differing area assignments. Our approach allowed us to examine how lumping and splitting both taxa and areas influenced the probability of inferring the ancestral origin of Polioptilidae and the point in the tree where taxa dispersed between North and South America.

## MATERIALS AND METHODS

### Taxon sampling and laboratory protocols

We sampled the 15 species in Polioptilidae currently recognized by the American Ornithological Society (Fig. 1; Chesser *et al.*, 2015; Remsen *et al.*, 2015). Our sampling also included 95% (Dickinson & Christidis, 2014; *n =* 61 ssp) or 98% (Clements *et al.*, 2016; *n* = 58 ssp) of all widely recognized subspecies, and where possible, sampling from throughout each species’ distribution (Table S1). We also included the most recently described taxa in the family, *P. clemensti* (Whitney & Alonso, 2005) and *P. guianensis attenboroughi* (Whittaker *et al.*, 2013). We lacked samples from four subspecies recognized by Dickinson & Christidis (2014), including *M. cinereiventris albapiculus* (Olson, 1980), *M. cinereiventris unicus* (Olson, 1980), *R. melanurus pallidus* (Todd, 1913), *P. caerulea comiteca* (Phillips, 1991), and a single subspecies recognized by Clements *et al.*, (2016) *P. caerulea gracilis* (van Rossem & Hachisuka, 1937). Subspecies IDs were based on specimen labels and/or geographic location of each specimen. In a few cases subspecies descriptions were vague or incomplete so we assigned those samples to taxa whose range was close by.

**Figure 1.**
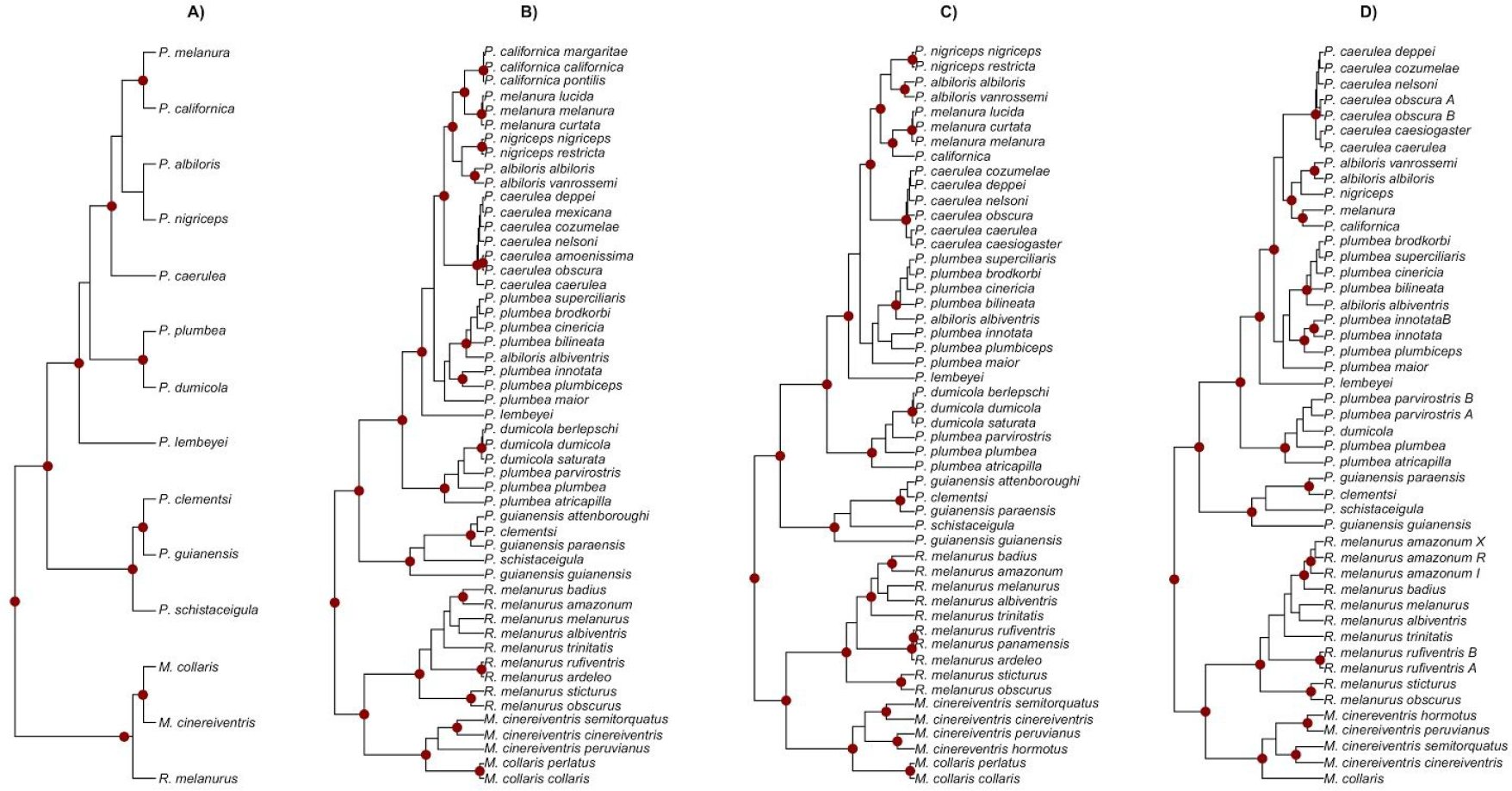
Maximum clade credibility species trees. Terminal tips in each of the species trees were currently recognized biological species (A), species and subspecies from Clements (B), species and subspecies from Howard & Moore (C), and statistically inferred species from BPP (D). Nodes with posterior probabilities ≥ 0.95 are denoted with red dots.

We extracted genomic DNA from tissue or museum skin toe-pads using the DNeasy tissue extraction kit (Qiagen, Valenica, CA). We collected the mitochondrial gene NADH: ubiquinone oxidoreductase core subunit 2 (ND2) from all samples (*n* = 333) and we downloaded ND2 sequences of *P. californica* (*n =* 38; Zink *et al.*, 2013) and *M. collaris* (*n* = 10; Naka *et al.*, 2012) from GenBank. We also collected six nuclear markers; Eukaryotic translation elongation factor 2 (EEF2; *n* = 113), muscle, skeletal, receptor tyrosine kinase (MUSK I-3; n = 97), Aconitase 1 (ACO1 I-9; *n* = 105), Beta fibrinogen Intron 5 (FIB I-5; *n* = 128), Myoglobin (MYO I-2; *n* = 103), and Ornithine decarboxylase (ODC I6-I7; *n* = 123), from all described taxa and select samples across species ranges for which fresh tissue was available. Sampled taxa lacking tissues (*M. cinereiventris magdalenae*, *M. collaris paraguensis, M. collaris torquatus, M. collaris colombianus, R. melanurus austerus*, *R. melanurus duidae*, *R. melanurus griseodorsalis*, *R. melanurus pallidus*, *R. melanurus sanctaemarthae*, *P. guianensis facilis, P. plumbea anteocularis, P. plumbea daguae, P. caerulea perplexa*, and *P. lactea*) were represented in our dataset by only ND2 sequences. For the ND2 dataset *Troglodytes aedon*, *T. brunneicollis*, *T. musculus*, *T. ochraceus*, *T. rufociliatus*, *T. rufulus*, and *T. solstitialis* were specified as the outgroups because of the sister relationship between Polioptilidae and Troglodytidae (Barker *et al.,* 2004). We amplified loci via polymerase chain reaction (PCR) in 12.5 μl reactions using the following protocol: denaturation at 94 °C for 10 min, 40 cycles of 94 °C for 30 s, using a gene specific annealing temperature (see below) for 45 s, and 72 °C for 2 min, followed by 10 min elongation at 72 °C and 4 °C soak. Locus specific annealing temperatures were as follows; ND2: 54 °C; EEF2, MUSK: 58 °C; ACO1, FIBI5, MYOI2: 60 °C; ODC: 65 °C. We used the nuclear primer sequences from Kimball *et al.*, (2009) and we designed internal primers for the ND2 gene to amplify DNA from museum skins. PCR products were sent to the High-Throughput Genomics Unit (University of Washington) for all subsequent steps. Briefly, the facility purified PCR products using ExoSAP-IT (USB Corporation, Cambridge, MA), performed cycle-sequencing reactions, and sequenced the final products using BigDye (Applied Biosystems, Foster City, CA) on a high-throughput capillary sequencer. We aligned chromatograms in Sequencher 4.9 (GeneCodes Corporation, Ann Arbor, MI) and Geneious v. 9 (Kearse *et al.*, 2012). We resolved nuclear markers that had insertion/deletion events between homologous nuclear alleles using the program Indelligent (Dmitriev & Rakitov, 2008). To estimate the gametic phase of heterozygous sites in the nuclear markers, we performed two separate runs for each marker in the program PHASE v. 1.2 (Stephens *et al.*, 2001) and we assigned sites that had posterior probabilities of < 0.90 as ambiguous. We aligned nuclear loci before and after phasing using MUSCLE in MEGA6 (Tamura *et al.*, 2013). Variation in sample size among nuclear loci reflects the relative degree of success in our polymerase chain reaction (PCR) amplifications. DNA sequences are available on GenBank (MG902958 - MG903296; Intron numbers pending).

### mtDNA tree estimation

We determined the best-fit sequence model for each gene alignment based on the Akaike Information Criterion (AIC) scores using MEGA6 (Tamura *et al.*, 2013). To estimate a gene tree for the complete ND2 dataset (*n* = 388; including *Troglodytes aedon*, *T. brunneicollis*, *T. musculus*, *T. ochraceus*, *T. rufociliatus*, *T. rufulus*, and *T. solstitialis* as outgroups), we ran MrBayes v. 3.2.5 (Ronquist & Huelsenbeck, 2003) for 15,000,000 generations with four runs and discarded the first 25% of the trees as the burn-in. We assessed MCMC convergence and determined burn-in by examining ESS values and likelihood plots in the program Tracer v. 1.5 (Rambaut & Drummond, 2010), and standard deviation of the split frequencies among runs in MrBayes. For the mtDNA tree the tip names follow the taxonomy of Clements *et al.* (2016).

### Species tree estimation

We estimated species trees from samples with both nuclear DNA and mtDNA using *BEAST (Heled & Drummond, 2010), a part of the BEAST v.2.2.1 package (Bouckaert *et al.*, 2014). To assign alleles to a taxon we used four alternative taxonomic treatments: 1) currently recognized biological species (*n* = 14; Chesser *et al.*, 2015; Remsen *et al.*, 2015); 2) subspecies and monotypic species based on the taxonomy of Clements *et al.* (2016; hereafter Clements; *n* = 51); 3) subspecies and monotypic species based on the Howard & Moore check-list (Dickinson & Christidis, 2014; *n* = 50); and 4) statistically inferred species using the Bayesian Phylogenetics & Phylogeography program (*n* = 46; see species delimitation section below). Because there are no known fossils within Polioptilidae, we time-calibrated the species tree using published substitution rates, assuming a generation time of one year based on the age when *P. caerulea* begins breeding (Kershner & Ellison, 2013). We specified lognormal distributions for the relaxed uncorrelated rates for all loci (ND2: mean = 0.0125; SD = 0.1; Smith & Klicka, 2010; autosomal markers: mean = 0.00135; SD = 0.45; Ellegren, 2007; sex-linked nuclear markers: ACO-1 I-9 and MUSK I-3: mean = 0.00145; SD = 0.45; Ellegren, 2007). We used mutation rates to obtain more accurate branch lengths but we do report divergence time estimates. Preliminary analyses determined that 1,000,000,000 generations were required to obtain high ESS values (sampling every 10,000 generations). The locus-specific substitution models were as follows: ND2: GTR + G; ACO-1 I-9: HKY + G; MUSK I-3: HKY + G; EEF2: K2 + G; FIB I-5: HKY +G; MYO I-2: K2 + G; ODC I6-I7: HKY + G. We specified a Yule process on the species tree prior and lognormal distributions on sequence model prior distributions. We followed the same MCMC diagnostics described above. Finally, we summarized the posterior distribution of trees by building a maximum credibility clade (MCC) tree in TreeAnnotator v2.2.1 (Bouckaert *et al.*, 2014). Visual plots of phylogenetic trees were produced using the APE (Paradis *et al.*, 2004) and ggtree (Yu *et al.*, 2017) R packages.

### Species delimitation

There are a number of software packages and approaches available for delimiting species from DNA sequence data (e.g., Reid *et al.*, 2012; Yang & Rannala, 2010; Jackson *et al.*, 2017). In this study we used the program Bayesian Phylogenetics & Phylogeography (BPP, v.3.2), which models multilocus sequence data to generate probabilities for lineages identified *a priori* as potential species-level taxa (Rannala & Yang, 2003; Yang & Rannala, 2010). The utility of BPP for delimiting species has been questioned (Sukumaran & Knowles, 2017), and others have advocated for using multiple approaches (Satler *et al.*, 2013; Carstens *et al.*, 2013), but the method remains a robust means of inferring species from multilocus data (Rannala, 2015; Leaché *et al.*, 2018).

BPP uses a Bayesian modeling approach to generate a posterior distribution of speciation models where the number of species in each model varies. Speciation probabilities are estimated from the sum of probabilities of all models for speciation events. The BPP model accounts for gene tree uncertainty and incomplete lineage sorting, and assumes no gene flow among species after divergence, no recombination among loci, and implements the JC69 (Jukes & Cantor, 1969) mutation model. BPP includes divergence time parameters (τ) and population size parameters (θ = 4N_e_μ), where N_e_ is the effective population size and μ is the mutation rate per site per generation. We ran the analyses using the A10 algorithm (species delimitation with user-specified guide tree) for 500,000 generations, sampling every five generations, and specified a burn-in of the first 25,000 generations. To assess the robustness of speciation probabilities, we implemented three combinations of priors that represented different population sizes and different ages for the root in the species tree: (i) large N_e_ and deep divergence: θ and τ gamma priors G(1, 10) and G(1, 10); (ii) small N_e_ and shallow divergence: θ and τ gamma priors G(2, 2000) and G(2, 2000), and (iii) large N_e_ and shallow divergence: θ and τ gamma priors G(1, 10) and G(2, 2000). We performed BPP analyses using the four guide trees (the topologies estimated in *BEAST) that represented our four alternative taxonomic treatments. To generate the guide tree for the statistically inferred taxa, we first estimated an additional species tree where we assigned alleles to lineages defined by geographic and genetically differentiated mtDNA groups (*n* = 49). In cases where species posterior probabilities < 0.95 among three taxa, we changed the topological relationships among the three subspecies involved to determine speciation probabilities under each possible arrangement. All analyses were done first using all loci (one mtDNA locus and six nuclear loci) and then again with only nuclear loci (*n* = 6). We considered terminal lineages from nodes to be statistically supported species when speciation posterior probabilities generated using the all-loci dataset were ≥ 0.95 under all three prior combinations (i.e., permutations on population size and root age) regardless of guide tree.

### Probabilistic modeling of geographic range evolution

To infer the geographic origins of Polioptilidae, we employed probabilistic modeling of geographic range evolution using the R (2015) package BioGeoBEARS (Matzke, 2014). We performed independent analyses using the four species trees generated using alternative taxonomic treatments. Terminal tips were assigned to geographic areas based on their sampled localities. We first assigned taxa to these two broad-scale geographic regions (Fig. 6A), and then subdivided these two areas first into four (Fig. 6B) and then into eight (Fig. 6C) geographic areas. The geographic areas used in each partition were as follows (Fig. 6, top panel, maps A, B, & C): *Two areas*: 1) North America - west of the Isthmus of Panama through Central America, Mexico, and the Nearctic region and Cuba. 2) South America - east of the Isthmus of Panama through the South American continent; *Four areas*: 1) Nearctic region - the United States of America, Canada, and northern Mexico, 2) Cuba, 3) Middle America - tropical lowland Mexico to the Isthmus of Panama, and 4) South America - east of the Isthmus of Panama through the South American continent. *Eight areas*: 1) Nearctic region - the United States of America, Canada, and northern Mexico, 2) Cuba, 3) Middle America - tropical lowland Mexico to the Isthmus of Panama, 4) west of the Andes - east of the Isthmus of Panama through the Darien, Choco, and Magdalena Valley, 5) Maranon Valley - an inter-Andean valley in northern Peru, 6) Amazon Basin - the lowland forests in the Amazon Basin, 7) Dry Diagonal - the arid region in eastern and northeastern Brazil, and 8) Atlantic Forest - the tropical rainforest biome that extends along the Atlantic coast of Brazil.

We used two historical biogeographic models, the Dispersal–Extinction–Cladogenesis (DEC) model (Ree & Smith, 2008) and the DEC+J model that allows for jump dispersal founder events (Matzke, 2014). We set the maximum range size to equal the maximum number of ranges occupied by a species in each treatment. The maximum number of allowed ranges equaled two areas in all cases except the biological species analysis with eight areas, where *P. plumbea* occurred in five of them. In the four-area analyses, we constrained ancestral areas to only adjacent areas that were consistent with species’ distributions. We performed AIC model selection by estimating the likelihood of the data under the DEC and DEC+J models, comparing the difference in AIC values, and calculating AIC weights. For ongoing debate about the DEC+J model and biogeographic model selection see Ree & Sanmartín (2018).

## RESULTS

### Species and mtDNA tree

The species trees inferred using each of our taxonomic treatments and the mtDNA tree recovered similar clades (Figs. 1-4). *Ramphocaenus* and *Microbates* formed a well-supported sister group. This clade was in turn sister to a monophyletic *Polioptila*, which was divided into two main clades, the *P. guianensis* complex and the remaining taxa (Fig. 1; Fig. S1). Support for nodes was generally high except within the clade containing North American taxa. The phylogenetic placement of *P. lactea* was only inferred from mtDNA (Fig. 4). Relationships among nodes common to the four-species trees (Fig. 1) and the mtDNA tree (Figs. 2-4: Fig. S1) were congruent except in the placement of *P. plumbea* (Fig. 1). The majority of deep nodes in the species tree were supported, but numerous relationships at the tips were unresolved. The species trees using subspecies and statistically inferred taxa showed that *P. plumbea* was not monophyletic (Fig. 1), and the lumping of *P. plumbea* samples in the biological species tree produced an alternative placement of the taxon with low support (Fig. 1). Hereafter, all topological relationships in this paragraph refer to the species trees with terminal units based on statistically inferred taxa, Clements, or Howard & Moore, unless stated otherwise. For clarity, we discuss subspecies according to Clements except where noted. The *Polioptila guianensis* complex (*P. guianensis, P. clementsi,* and *P. schistaceigula)* was sister to all other taxa in the genus. Within the remaining *Polioptila,* the first divergence was between a clade composed of a subset of *P. plumbea* subspecies (*atricapilla*, *plumbea*, *parvirostris*) and *P. dumicola*. In the mtDNA tree, *P. dumicola* was sister to this clade of *P. plumbea* (Fig. 4). The next node separated the Cuban endemic *P. lembeyei* from the remaining *Polioptila*. These included the remaining *P. plumbea subspecies* (*maior*, *innotata, plumbiceps*, *bilineata*, *cinceria*, *superciliaris*, and *brodkorbi*), which were sister, albeit with low support, to a clade containing the more northerly distributed forms *P. californica*, *P. melanura*, *P. nigriceps*, *P. albiloris,* and *P. caerulea* (Figs. 1 & 4; Fig. S1). Several species, particularly the more widely distributed ones, showed deep phylogeographic structure largely concordant with subspecific variation (discussed below), but *P. plumbea*, *P. albiloris* and *P. guianensis* were not monophyletic.

**Figure 2.**
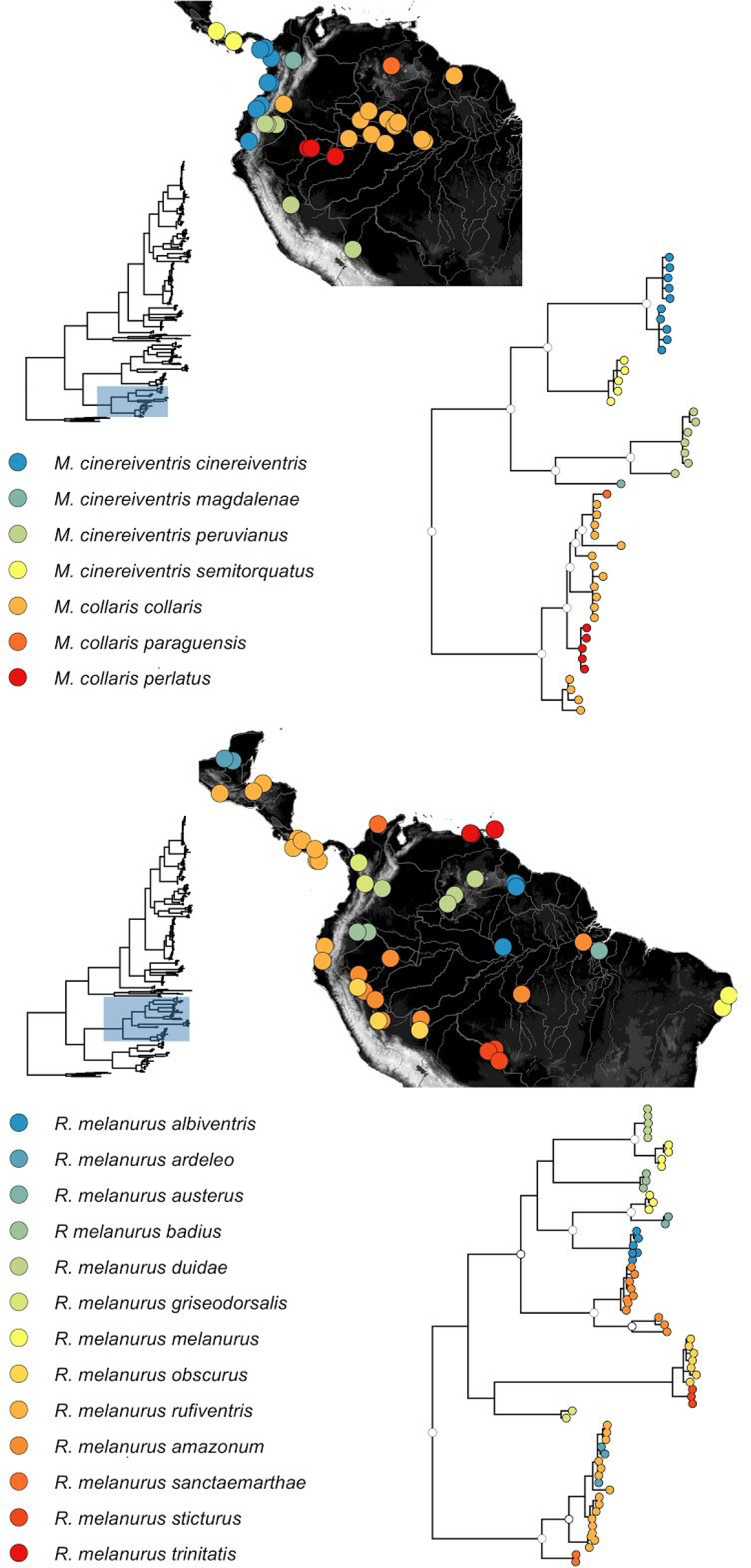
mtDNA trees for *Microbates* and *Ramphocaenus* and sampling maps. Samples are colored coded according to subspecies designation of Clements. White nodes have posterior probabilities of 0.95.

### Species limits

*Microbates*. – Subspecies in *M. cinereiventris* formed geographically and genetically distinct clades, although *M. cinereiventris peruvianus* and *M. cinereiventris magdalena* were represented by single exemplars. Only one of these (*M. cinereiventris peruvianus*) was represented with multilocus data (Figs. 1 & 2). *Microbates cinereiventris cinereiventris* (east Panama through the Choco) and *M. cinereiventris semitorquatus* (western Panama through Costa Rica) were sister taxa and formed a clade distributed west of the Andes (Figs. 1 & 2; Fig. S1). *Microbates cinereiventris peruvianus and M. cinereiventris hormotus* were not reciprocally monophyletic, but the *M. cinereiventris peruvianus* sample from Bolivia is genetically differentiated from the others. *Microbates collaris* showed slight genetic structuring across northern Amazonia, but this variation did not correspond neatly with subspecies (*collaris*, *colombianus*, *paraguensis*, and *perlatus*) boundaries (Fig. 2; Fig. S1). Based on the BPP analyses, in *Microbates* 5–6 species were inferred: Clements (PP = 0.997, *n* = 5), Howard & Moore (PP = 0.986, *n* = 6), and phylotaxa (PP = 0.986, *n* = 6; Fig. 5). The splitting of *M. collaris collaris* and *M. c. perlatus* was not robust to the prior or the exclusion of the mtDNA, the posterior probabilities from all other speciation events in *Microbates* were (Fig. 5).

*Ramphocaenus*. – The majority of taxa were nested within geographically coherent clades and were reciprocally monophyletic (Fig. 1; Fig. 2; Fig. S1). A clade containing *Ramphocaenus melanurus sticturus* and *R. melanurus obscurus* was sister to all other *R. melanurus* taxa, and a subsequent clade composed of the taxa occurring west of the Andes was sister to all taxa from east of the Andes (Fig. 1; Fig. 2; Fig. S1). The mtDNA tree included the Magdalena Valley endemic *R. melanurus griseodorsalis*, but its position was unresolved (Fig. 2; Fig. S1). Also unique to the mtDNA tree was the Santa Marta Mountain endemic *R. melanurus sanctaemarthae*, which was nested within a clade containing the taxa from Central America, coastal Colombia, and Ecuador (Choco) (Fig. 2; Fig. S1). The subspecies *Ramphocaenus melanurus rufiventris*, *R. melanurus ardeleo*, and *R. melanurus panamensis* were not genetically sorted (Fig. S1). BPP analyses identified 9–10 species with a posterior probability > 0.95 that were robust to the prior across the three specified taxonomic treatments. The models with the highest posterior probabilities for each taxonomic treatment group were as follows: Clements (PP = 0.997, *n* = 9), Howard & Moore (PP = 0.606, *n* = 9), and phylotaxa (PP = 0.941, *n* = 11; Fig. 5).

With the exception of the *R. m. panamensis* (Howard & Moore), all taxa in the BPP analysis for *Ramphocaenus* had high posterior probabilities (> 0.99; see Table S2). The BPP analyses based on phylotaxa found support for undescribed lineages within *R. melanurus rufiventris* and *R. melanurus amazonum* with the all loci dataset, but when mtDNA were excluded, the posterior probabilities for these splits were lower (0.5–0.8; Fig. 5). Individuals of *R. melanurus rufiventris* from Quetzaltenango, Guatemala were differentiated from all other *R. melanurus rufiventris* (Yucatan, Mexico to Tumbes, Peru). Within *R. melanurus amazonum* there were three generally well-supported lineages (PP = 0.941–1.0) that correspond to the Inambari, Rondonia, and Xingu endemic areas. More precise geographical distributions for these putative taxa are unclear because our sampling was incomplete across southern Amazonia.

*Polioptila guianensis* complex. – *Polioptila schistaceigula* and *P. clemensti* were nested within *P. guianensis* (Figs. 1 & Figs. 3; Fig. S1). *Polioptila guianensis facilis* and *P. guianensis paraensis* formed a clade with *P. clementsi* and the recently described *P. guianensis attenboroughi*. In the species tree and mtDNA tree, the relationships among *P. guianensis paraensis, P. guianensis attenboroughi* and *P. clementsi* were unresolved (Fig. 3; Fig. S1). BPP estimated that three of the five taxa (*P. guianensis facilis* was exclude because of a lack of nuclear DNA) in this clade had speciation probabilities > 0.95 under all three priors (PP = 0.642, *n* = 4; Fig. 5), and similar results were obtained with datasets containing all loci and only nuclear DNA loci (Fig. 5). Speciation probabilities for the splitting of *P. clementsi* and *P. guianensis attenboroughi* ranged from 0.349 to 0.810, and these were the only taxa in this complex with speciation probabilities lower than the acceptance threshold (Fig. 5).

**Figure 3.**
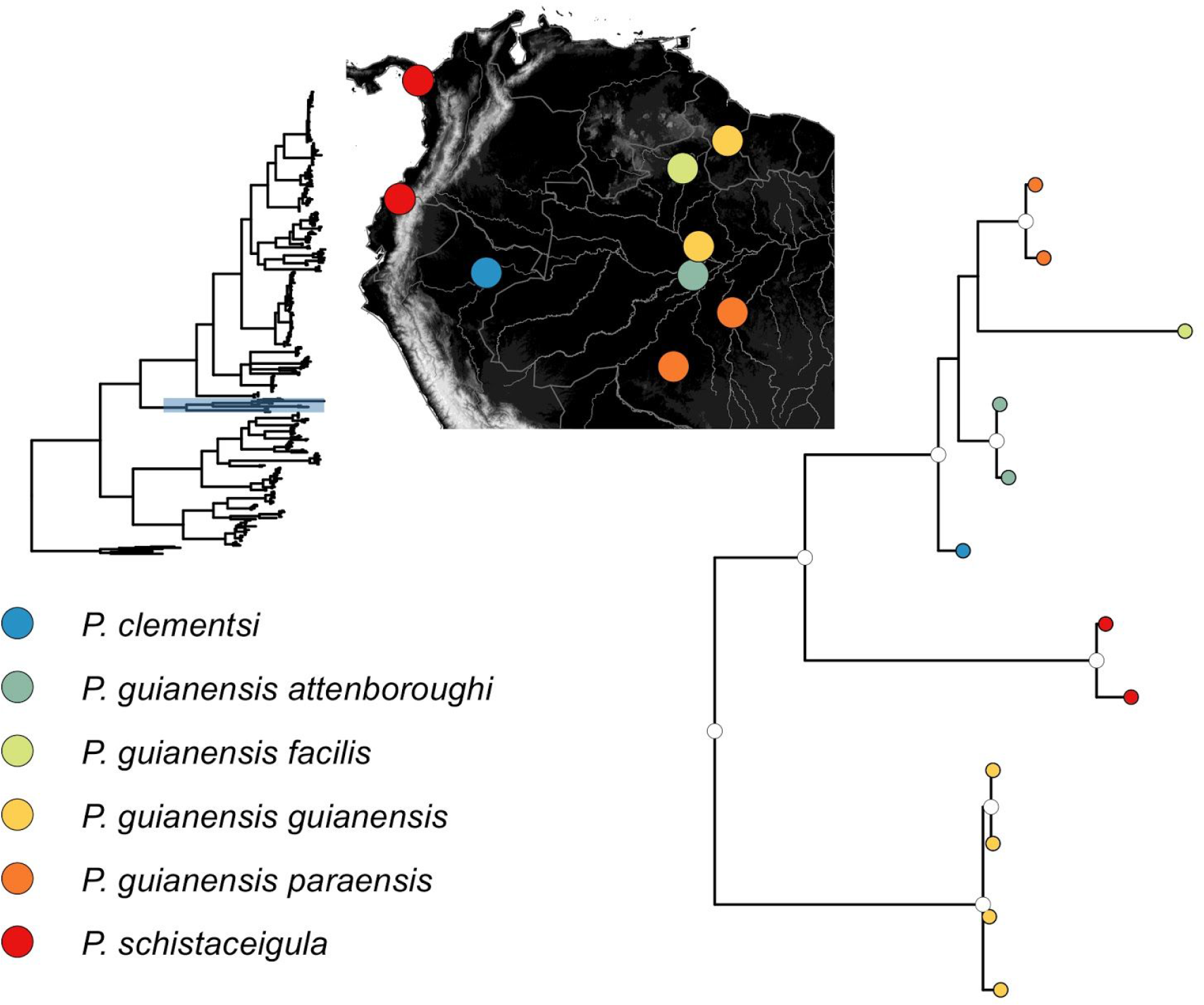
mtDNA trees for *Polioptila guianensis* complex and sampling maps. Samples are colored coded according to subspecies designation of Clements. White nodes have posterior probabilities of 0.95.

*Polioptila*. – The remaining *Polioptila* taxa harbored considerable geographically structured genetic diversity. *Polioptila albiloris* showed a phylogeographic break that matches the subspecific break across the Isthmus of Tehuantepec (Figs. 1 & 4; Fig. S1). Nominate *P. albiloris* occurred east of the Isthmus, from El Salvador through Costa Rica (Figs. 1 & 4; Fig. S1), whereas *P. albiloris vanrossemi* was distributed west of the divide, along the Pacific coast of Mexico (Figs. 1 & 4; Fig. S1). The third taxon in *P. albiloris, P. a. albiventris*, was nested within *P. plumbea*, rendering *P. albiloris* paraphyletic. *Polioptila caerulea* showed deep phylogeographic structure somewhat concordant with morphologically defined subspecies. mtDNA haplotypes in *P. caerulea nelsoni*, *P. caerulea cozumelae*, and *P. caerulea caerulea* were geographically sorted, but the validity of *P. caerulea deppei* was less certain as haplotypes were grouped with those representing *P. caerulea amoenissima* and *P. caerulea mexicana* (Fig. 4; Fig. S1). The single sample of *P. caerulea caesiogaster* (not recognized by Clements) from the Bahamas also had a unique haplotype. Two additional groups identified represented a split between western coastal (*P. caerulea amoenissima* A) and more interior *P. caerulea* populations (*P. caerulea amoenissima* B), although these forms appear to co-occur in southern Nevada. This mtDNA divide is not suggested by either the Clements nor Howard & Moore taxonomies. *Polioptila plumbea* formed two clades that corresponded well with geography. One clade included all *P. plumbea* taxa distributed in Central America (*brodkorbi, cinericia, superciliaris*) west of the Andes (*daguae, bilineata*), the Maranon Valley in eastern Peru (*maior*), and throughout northern South America (*anteocularis, plumbiceps, innotata*). Surprisingly, this clade also includes a morphological outlier, the Yucatan Peninsula form *P. albiloris albiventris* (Figs. 1 & 4; Fig. S1). The remaining *P. plumbea* subspecies occur east of the Andes (*parvirostris, atricapilla, plumbea*), comprising a second clade that is more closely related to the southern South American forms *P. lactea* and *P. dumicola*. All *P. plumbea* taxa were reciprocally monophyletic in the mtDNA tree (Fig. 4; Fig. S1). The western Amazonian *P. plumbea parvirostris* was divergent across the Maranon and Ucayali rivers. Additional taxa that were not reciprocally monophyletic in the mtDNA tree were *P. caerulea* (*perplexa, mexicana,* and *deppei*)*, P. melanura* (*melanura, lucida, and curtata*), *P. californica* (*californica, margaritae,* and *pontilis*), *P. nigriceps (nigriceps and restricta),* and *P. dumicola* (*dumicola, berlepschi,* and *saturata*) (Fig. 4; Fig. S1). Sample sizes in *P. lactea* (*n* = 3) were too small to assess phylogeographic variation.

**Figure 4.**
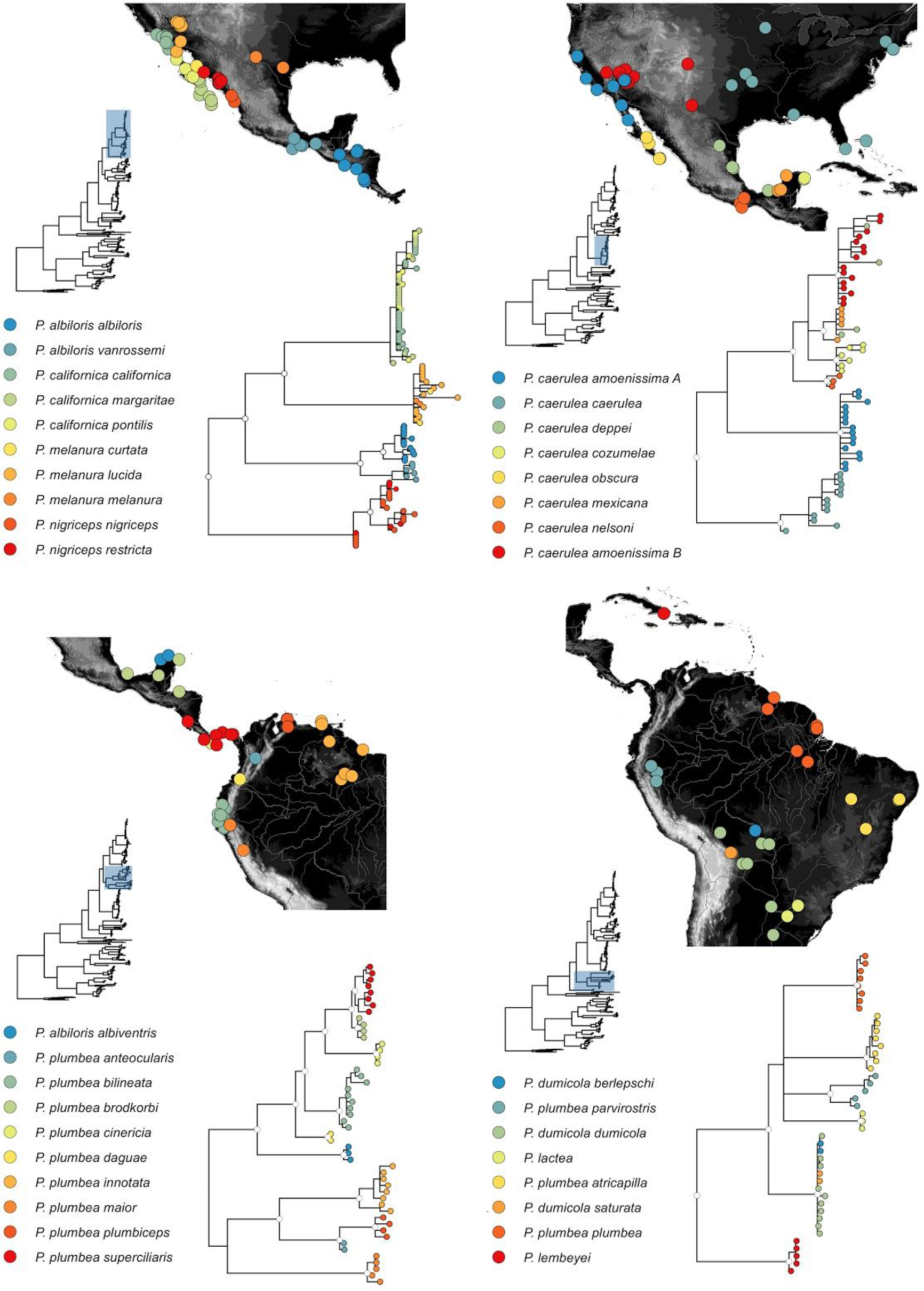
mtDNA trees for *Polioptila* clades and sampling maps. Samples are colored coded according to subspecies designation of Clements. White nodes have posterior probabilities of 0.95.

**Figure 5.**
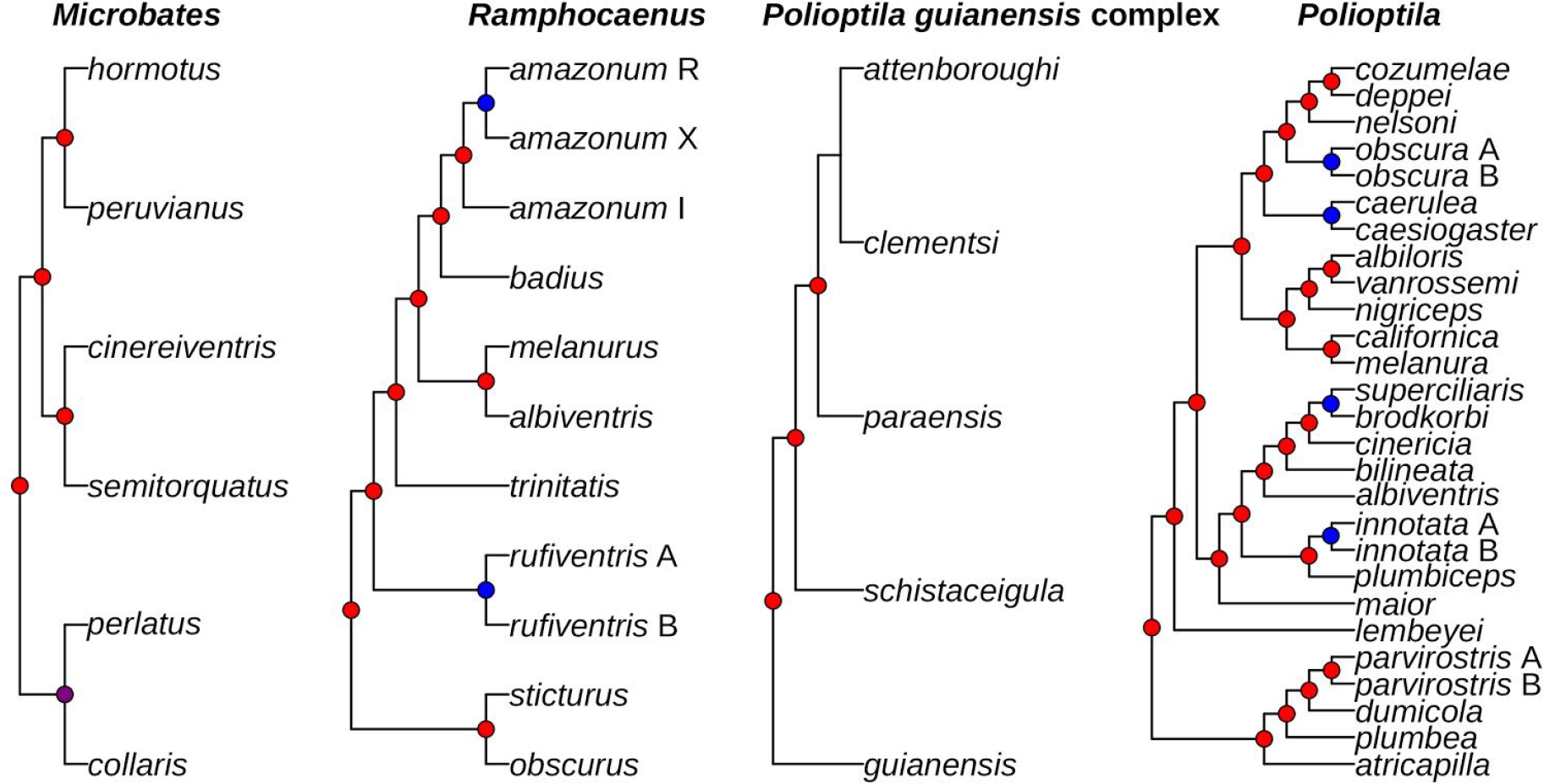
Stability of speciation probabilities across priors and datasets. Shown are topologies for *Microbates*, *Ramphocaenus*, the *P. guianensis* complex, and *Polioptila* (sans, *P. guianensis* and allies) using the phylotaxa taxonomy. Colored nodes represent speciation posterior probabilities values. Shown are colored nodes for speciation events that are stable across all priors and datasets (red), unsupported when mtDNA was removed (purple), unstable across priors and datasets (blue), and never supported (no node).

**Figure 6.**
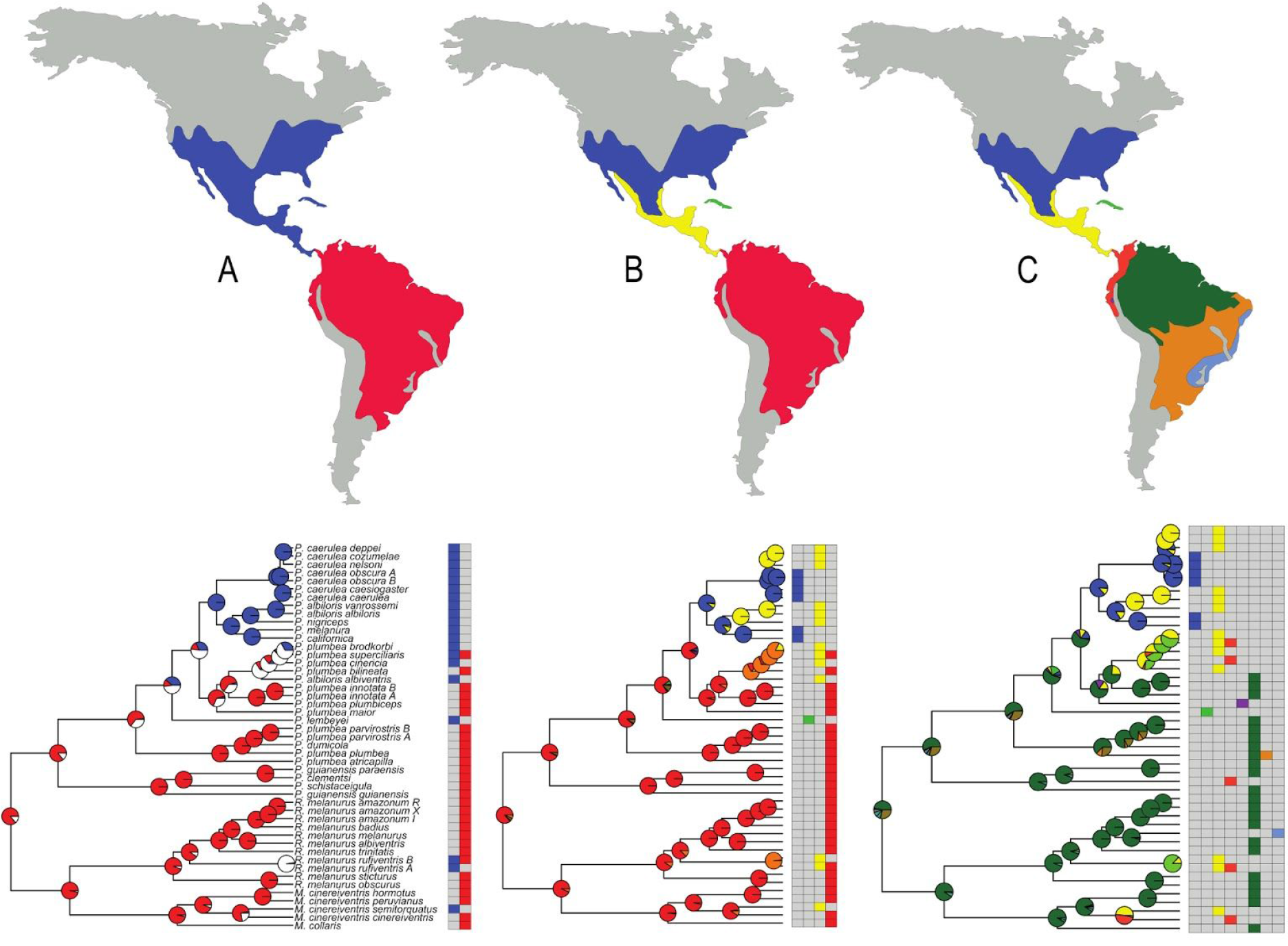
Alternative biogeographic areas for the New World and probabilistic biogeographic modeling on the species trees using statistically inferred taxa. The New World was geographically partitioned into two (A), four (B), or eight (C) areas. The geographic areas used in each partition were as follows (Fig. 2, top panel, maps A, B, & C): *Two areas*: 1) North America - west of the Isthmus of Panama through Central America, Mexico, and the Nearctic region and Cuba. 2) South America - east of the Isthmus of Panama through the South American continent; *Four areas*: 1) Nearctic region - the United States of America, Canada, and northern Mexico, 2) Cuba, 3) Middle America - tropical lowland Mexico to the Isthmus of Panama, and 4) South America - east of the Isthmus of Panama through the South American continent. *Eight areas*: 1) Nearctic region - the United States of America, Canada, and northern Mexico, 2) Cuba, 3) Middle America - tropical lowland Mexico to the Isthmus of Panama, 4) west of the Andes - east of the Isthmus of Panama through the Darien, Choco, and Magdalena Valley, 5) Maranon Valley - an inter-Andean valley in northern Peru, 6) Amazon Basin - the lowland forests in the Amazon Basin, 7) Dry Diagonal - the arid region in eastern and northeastern Brazil, and 8) Atlantic Forest - the tropical rainforest biome that extends along the Atlantic coast of Brazil.

The number of putative *Polioptila* species Clements (*n* = 31), Howard & Moore (*n* = 29), and phylotaxa (*n* = 27) identified using the BPP models based on the all loci dataset varied across taxonomic treatments. The models with the highest posterior probabilities for each treatment were as follows: Clements (PP = 0.277, n = 29), Howard & Moore (PP = 0.522, *n* = 28), and phylotaxa (PP = 0.972, n = 27; Fig. 5). We obtained similar results when we excluded the mtDNA locus but speciation probabilities were lower for some splits. For the phylotaxa treatment, we found support (PP > 0.95) for distinct lineages within *P. caerulea obscura*, *P. plumbea innotata*, and *P. plumbea parvirostris* using the all loci dataset, but not with only the nuclear DNA dataset (Fig. 5). Based on our sampling, the break in *P. plumbea parvirostris* corresponded to north and south of the Maranon River, and in *P. plumbea innotata* the break was north and south of the Tepuis in Venezuela. Support for the east-west split in *P. caerulea obscura* ranged from 0.080–1.0 across the priors when the mtDNA was excluded. A similar result was observed for the alternative treatment using Clements, where the putative species *P. caerulea amoenissima* and *P. caerulea obscura* had high speciation probabilities (> 0.95) with the all loci dataset, but low probabilities (< 0.5) with only nuclear loci. The split between *P. plumbea brodkorbi* and *P. plumbea superciliaris* also had lower speciation probabilities when mtDNA was excluded (Fig. 5). Support for the differentiation of subspecies within *P. californica*, *P. melanura*, *P. nigriceps,* and *P. dumicola* were not robust to the prior and the guide tree. All other putative species in *Polioptila* species tested in our analysis had consistently high posterior probabilities.

### Biogeographic modeling

All of our BioGeoBEARS analyses suggest that Polioptilidae evolved in South America with multiple colonizations of North America and a single dispersal event into Cuba (Table 1; Fig. 6; Figs. S2-S4). The probability of ancestral areas was influenced by the biogeographic model, the number of areas, and the species tree. The data were more likely when we used the DEC+J model than the DEC model in 9 out of 12 cases (Table 2). The change in AIC values between the DEC and DEC+J models was typically greater when the number of areas increased (two to eight) and when the tree had a greater number of terminal tips (Table 1). When the number of areas was two, only the species tree using statistically inferred taxa had a higher AIC score for the DEC+J model over the DEC model (Table 1). For the basal node of the Polioptilidae tree, we summed probabilities into bins representing North or South American origin, or ambiguous. South America was the most probable (0.59–0.95) ancestral area at this node in 11/12 tests, and the probability of a North American origin was less than 0.02 in all cases (Table 1). The lone exception was when we used eight areas and the biological species tree, for which ambiguous had the highest probability (0.55) for the ancestral area at the base of Polioptilidae (Table 1). A South American ancestral origin was most likely when we used the subspecies and using statistically inferred taxa species trees (Table 1). Long-distance migration, present in the *P. caerulea* subspecies that breed in eastern and western North America, appears to have evolved relatively recently and after the ancestor had dispersed northward.

**Table 1.** Probability of ancestral origin of Polioptilidae using alternative taxonomies and biogeographic area assignments. Shown are the summed probabilities for Polioptilidae originating in in only North or only South America, or in both (Ambiguous).

| **Dataset** | **North**<br>**American** | **South**<br>**American** | **Ambiguous** |
| --- | --- | --- | --- |
| **Biological Species** |  |  |  |
| Two Areas | 0.01 | 0.61 | 0.38 |
| Four Areas | 0.07 | 0.59 | 0.34 |
| Eight Areas | 0.02 | 0.44 | 0.55 |
| **Clements** |  |  |  |
| Two Areas | 0.00 | 0.86 | 0.14 |
| Four Areas | 0.00 | 0.95 | 0.05 |
| Eight Areas | 0.00 | 0.77 | 0.23 |
| **Howard & Moore** |  |  |  |
| Two Areas | 0.00 | 0.83 | 0.17 |
| Four Areas | 0.00 | 0.92 | 0.08 |
| Eight Areas | 0.00 | 0.80 | 0.20 |
| **BPP** |  |  |  |
| Two Areas | 0.00 | 0.87 | 0.13 |
| Four Areas | 0.00 | 0.94 | 0.06 |
| Eight Areas | 0.00 | 0.84 | 0.16 |

**Table 2.** Biogeographic model selection. Shown are each taxonomic partition, the number of total biogeographic areas used in the analyses, AIC score for the DEC and DEC+J models, and AIC weights (AIC wt) for each model.

| **Taxonomy** | **Number of Areas** | **AIC DEC** | **AIC DEC+J** | **AIC wt DEC** | **AIC wt DEC+J** |
| --- | --- | --- | --- | --- | --- |
| Biological Species | 2 | 24.65 | 26.65 | 0.73 | 0.27 |
|  | 4 | 48.94 | 45.88 | 0.18 | 0.82 |
|  | 8 | 91.37 | 88.3 | 0.18 | 0.82 |
| Clements | 2 | 52.66 | 52.57 | 0.49 | 0.51 |
|  | 4 | 89.22 | 81.05 | 0.02 | 0.98 |
|  | 8 | 170.6 | 144.1 | 0 | 1 |
| Howard & Moore | 2 | 60.98 | 62.7 | 0.7 | 0.3 |
|  | 4 | 96.61 | 91.82 | 0.08 | 0.92 |
|  | 8 | 172.7 | 155.9 | 0 | 1 |
| BPP | 2 | 54.55 | 46.09 | 0.01 | 0.99 |
|  | 4 | 91.68 | 75.34 | 0 | 1 |
|  | 8 | 164.8 | 131.1 | 0 | 1 |

## DISCUSSION

Species-tree and species-delimitation analyses provide a more nuanced and complete picture of the evolutionary history of Polioptilidae. The family is composed of three genera divided into four distinct groups. Currently recognized gnatcatchers and gnatwrens frequently harbor deep phylogeographic variation. Using coalescent modeling we inferred nearly three times more species than are currently recognized, and the majority of this diversity occurs in the Neotropics. The findings of our study represent a minimum diversity estimate because there are 19 taxa (one species, 18 subspecies) lacking multilocus data that were not included in our BPP analysis. Of the newly inferred species, 87% are concordant with taxa (monotypic species/subspecies) previously described based on subtle differences in plumage and size. In contrast, a relatively conserved morphology overall is at least in part responsible for apparent polyphyly inferred in three of the 14 species studied. Whether this reflects a retention of ancestral plumage characters in these groups or convergent evolution within more distantly related lineages is not clear. Our phylogeny suggests that the former is likely the case for *P. guianensis*, and the latter likely the case for *P. plumbea* and *P. albiloris*. All of the statistically inferred taxa should be considered fundamental evolutionary units for conservation decisions and evolutionary studies. For most, determining whether or not they are biological species will require additional work in genetic sampling, study of song, morphology, behavior, and putative hybrid zones.

Objectively defining the fundamental units of evolution (i.e., species) has important implications for historical biogeographic analysis. Probabilistic modeling of geographic ranges on the alternative species trees indicated that Polioptilidae probably had an ancestral origin in South America, with all three genera independently colonizing North America. Similar analyses on the biological species tree yielded a less certain biogeographic history. Our study corroborates the pervasive uncertainty around species limits expected in tropical clades and highlights the importance of using taxon units that accurately characterize diversity in biogeographic analyses.

### Delimiting diversity

Molecular systematic studies on Neotropical birds frequently recover paraphyletic species and deep divergences comparable to species-level differentiation (e.g., Chaves et al., 2011; Gutiérrez-Pinto *et al.*, 2012; Chaves et al., 2013; Smith *et al.*, 2013; Rheindt *et al.*, 2013; Cerqueira *et al.*, 2016; Ferreira *et al.,* 2017; Smith *et al.*, 2017; Musher *et al.*, 2018). In our study, we used species delimitation via coalescent modeling for an entire avian family and found that the majority of currently defined subspecies in Polioptilidae could be recognized as species based on their genealogical histories. Although species delimitation using the software BPP may be biased towards splitting in cases of high gene flow (Jackson *et al.*, 2017) the majority of the inferred species in our analyses have taxonomic names, which provides corroborating evidence the taxa are independently evolving units. Our results strongly contrast with a recent phenetics-based assessment of some taxa in the family (del Hoyo & Collar, 2017) that recognized only 15 species while retaining (or creating) paraphyletic species. Formal elevation of the statistically inferred taxa to biological species will likely await a more detailed understanding of the variation in song across currently recognized species’ ranges. In cases where songs have been compared among subspecies there is strong evidence for song differentiation. For example, *R. melanurus obscurus* (along with *R. melanurus sticturus)* is vocally distinct from that of parapatric forms *R. melanurus badius* and *R. melanurus amazonum* in western Amazonia (Harvey *et al.*, 2014b). Vocal differentiation among these taxa is consistent with the species tree topology that shows that *R. m. obscurus* and *R. m. sticturus* are sister to all other taxa in the clade. Most of the species identified in this paper are quantifiably different in phenotype or song. Rates of morphological change vary among taxonomic groups, and in some, phenotypic evolution and genetic evolution are decoupled (e.g., *Empidonax* flycatchers, Johnson & Cicero, 2002; Plethodon salamanders, Kozak *et al.*, 2009). In such groups, forms that appear very similar may represent lineages that have been evolving independently of one another for hundreds of thousands, perhaps millions, of years. A growing body of work suggests that such cryptic species are not evenly distributed, but rather, they are more likely to be uncovered in tropical latitudes (Tobias *et al.*, 2008; Barrowclough *et al.*, 2016). Species delimitation using the multispecies coalescent thus provides a powerful tool in categorizing biodiversity in such regions when traditional taxonomic methods fall short.

The non-monophyly of *P. plumbea, P. guianensis*, and *P. albiloris* pose immediate taxonomic problems. *Polioptila plumbea* is polyphyletic, forming two clades that are not sister, and both of these are paraphyletic. *Polioptila lactea* (mtDNA only, see Fig. 4) and *P. dumicola* are embedded within the *P. plumbea plumbea* “group” and *P. albiloris albiventris* is embedded within the *P. plumbea bilineata* “group”. *Polioptila plumbea brodkorbi* and *P. albiloris albiventris* do not appear to come into contact on the northern end of Yucatan Peninsula and both had high probabilities of being species based on the BPP analyses. To maintain monophyletic species or natural evolutionary groups, the majority of *P. plumbea* subspecies (and *P. albiloris albiventris*) should be elevated to full species. Such a change would be consistent with our results, as every *P. plumbea* subspecies analyzed with multilocus data was identified as a species-level taxon. Finally, *P. guianensis* is paraphyletic with respect to *P. schistaceigula* and *P. clementsi*. A conservative solution to this problem would be to divide the *P. guianensis* complex into three (minimally) taxa: 1) the nominate form *P. g. guianensis*; 2) *P. schistaceigula*; and 3) the remaining elements of the clade (*P. g. facilis*, *P. g. paraensis*, *P. guianensis attenboroughi*, and *P. clementsi* (see Fig. 1). Nevertheless, plumage differences coupled with genetic divergences among *P. g. facilis*, *P. g. paraensis*, *P. guianensis attenboroughi*, and *P. clementsi* (Whittaker *et al.*, 2013), suggest that this clade may be composed of more than one species.

A correlation between plumage differences (i.e., subspecies) and population history (i.e., genetic structure) was not consistently observed. In some examples described taxa showed genetic differentiation, but the depth of divergence, given our sampling, was not enough to be deemed species, based on coalescent modeling. This was the case for *P. guianensis attenboroughi* and *P. clementsi*, which had 1.4% uncorrected sequence divergence in mtDNA but were not strongly supported as separate species in the BPP species delimitation modeling. Among the strongest discrepancies between plumage differences and genetic differences were within *P. dumicola*, whose distinctive subspecies range from dark gray (*P. d. saturata*) to a much paler breast and belly (*P. d. berlepschi*). The lack of genetic depth among these taxa could be due to introgression, as the nominate form and *P. d. berlepschi* have come into increased contact as forest barriers have been removed (Atwood & Lerman, 2006). However, our sample sizes were too limited to evaluate potential hybridization between these forms. *Polioptila californica* subspecies were also not supported by species delimitation modeling, consistent with previous findings assessing depth of genetic variation across the species range (Zink *et al.*, 2013). In *P. californica*, our model results generally found low support for differentiation and the posterior probabilities were sensitive to alternative priors and guide trees. These taxon boundaries are among the most controversial in Polioptilidae (see Zink *et al.*, 2013; Zink *et al.*, 2016; McCormack & Maley, 2015). The failure of some taxa to be delimited using coalescent modeling could be due to a number of issues, including inadequate sampling of individuals and genetic variation, an insufficient number of genetic markers, rapid morphological evolution, or because some subspecies simply do not represent historically isolated lineages. Alternatively, existing coalescent methods may not be an appropriate for delimiting recently isolated taxa that experience continued gene flow, particularly when small multilocus datatsets are employed. Irrespective of these limitations, our findings provide a comparative framework for studying the evolutionary origins of taxa across the entire family.

### The influence of alternative taxonomies on biogeographic modeling

Using biological species as terminal units in the ancestral range models decreased our resolution of biogeographic history relative to that obtained using species trees based on subspecies or statistically inferred taxa. The majority of our analyses indicated there was a high probability that Polioptilidae originated in South America, but these probabilities were sensitive to how both species and areas were defined. Several of the major spatial-divergence events within Polioptilidae happened within species, thus lumped taxa and areas may not accurately designate at which node these events occurred. For example, when we used the four-area tree based on statistically inferred taxa (46 species), there was a high probability that *Microbates* and *Ramphocaenus* dispersed across the Isthmus of Panama at the node separating *M*. *semitorquatus* and *R. rufiventris* from their respective sister South American lineages. In contrast, when we used the four-area biological species tree (14 species), it was unclear when *Microbates* and *Ramphocaenus* dispersed into North America as there were similar probabilities of these events occurring at multiple nodes. Poor resolution of when these events occurred also decreased the probability of inferring with certainty the family’s continent of origin.

Polioptilidae is among the few New World avian families that probably originated in South America and yet were able to colonize temperate latitudes in North America (Smith *et al.*, 2012). Although support for our hypothesis is strong, our results are unable to account for extinct “ghost” lineages (Marshall, 2017) nor dynamic range shifts during glacial cycles (e.g., Zink & Gardner, 2017) that could change our interpretations. The inferred biogeographic pattern in Polioptilidae is consistent with the tropical niche conservatism hypothesis (Wiens & Donoghue, 2004) because the family has a higher tropical than temperate presence, and only one recent lineage has dispersed into temperate latitudes. Most New World tetrapod families inhabiting both tropical and temperate regions have a northern ancestry (Smith *et al.*, 2012). This ultimately may be the case for Polioptilidae as it may be secondarily South American because it is sister to the Troglodytidae (Barker *et al.*, 2004), and their common ancestor presumably colonized the New World from the Palaearctic via Beringia. The divergence of Polioptilidae and Troglodytidae may have arisen as a consequence of colonizing South America. Other clades of presumed tropical northern (e.g., Trogonidae, Momotidae) or secondarily South American (e.g., Thraupidae) origins do not currently occur in high northern temperate latitudes. Polioptilidae was likely able to expand past the Neotropical-Nearctic transition zone for two reasons. First, *Polioptila* species frequently occur in open and arid habitats so they are not subject to the desert filters between the Neotropical-Nearctic transition zone that likely constrain (in part) the northward expansion of most humid adapted tropical lineages (Ricklefs, 2002). Second, a few subspecies within *P. caerulea*, the northernmost lineage in the family, undergo partial or full seasonal migration. These characteristics are shared among the few other South American lineages (e.g., Tyrannidae; Trochilidae) that were able to colonize the Nearctic. In New World oscine songbirds (nine-primaried oscines), the most speciose clade of birds in the western hemisphere (Barker *et al.*, 2015), migration may have been achieved through evolutionary range shifts from temperate regions to more southern latitudes for the non-breeding winter season (Winger *et al.*, 2011; Winger *et al.*, 2014). In contrast, our analyses of Polioptilidae indicate that migratory behavior evolved very recently and from a tropical ancestor that expanded northward to breed in the temperate region of North America.

## SUPPLEMENTARY FIGURE LEGENDS (Supplementary material not included in this submission)

**Figure S1. mtDNA tree for Polioptilidae with zoomed-in subtrees.** Nodes with a posterior probability of 0.95 or higher are denoted with red circles. Names highlighted in bold are cases where the taxon has a different name between the Clements and Howard & Moore taxonomies, except for *Polioptila californica*. Howard & Moore does not recognize subspecies in *P. californica* so only the Clements taxonomy is shown for that taxon. In all other cases the first name shown is the Clements taxonomy and the Howard & Moore subspecies name is after the underscore.

**Figure S2-S4. Probabilistic modeling of ancestral areas for the biological species, Clements, and Howard & Moore taxonomic treatments (in order).** Shown are analyses run with two, four and eight biogeographical areas. *Two areas*: 1) North America - west of the Isthmus of Panama through Central America, Mexico, and the Nearctic region and Cuba. 2) South America - east of the Isthmus of Panama through the South American continent; *Four areas*: 1) Nearctic region - the United States of America, Canada, and northern Mexico, 2) Cuba, 3) Middle America - tropical lowland Mexico to the Isthmus of Panama, and 4) South America - east of the Isthmus of Panama through the South American continent. *Eight areas*: 1) Nearctic region - the United States of America, Canada, and northern Mexico, 2) Cuba, 3) Middle America - tropical lowland Mexico to the Isthmus of Panama, 4) west of the Andes - east of the Isthmus of Panama through the Darien, Choco, and Magdalena Valley, 5) Maranon Valley - an inter-Andean valley in northern Peru, 6) Amazon Basin - the lowland forests in the Amazon Basin, 7) Dry Diagonal - the arid region in eastern and northeastern Brazil, and 8) Atlantic Forest - the tropical rainforest biome that extends along the Atlantic coast of Brazil.

**Table S1. Taxon sampling, localities, and gene sampling per individual.**

## ACKNOWLEDGMENTS

We thank the following individuals and institutions that provided samples and support: N. Rice (ANSP), J. Dean (NMNH), R. Brumfield, F. Sheldon, D. Dittmann, S. Cardiff, V. Remsen (LSUMZ), D. Willard, S. Hackett, B. Marks (FMNH), J. Perez-Eman, (UCV & COP), M. Miller (STRI), P. Sweet, T. Trombone (AMNH), S. Birks (UWBM), P. Sykes (USGS), J. Trimble (MCZ), P. Escalante (CNAV), B. Hernandez-Banos (MCFZ), ZMUC (J. Fjeldså), LGEMA-IB-USP, and MMNH. We also thank L. Naka, G. Barrowclough, D. Lane, A. Cuervo, M. Harvey, R. Zink, B. Whitney, K. Provost, L. Musher, L. R. Moreira, J. Merwin, J. McCormack, and C. Smith. A. Alexio is supported by CNPq (#306843/2016-1).

